# Increased TRPV4 channel expression enhances and impairs blood vessel function in hypertension

**DOI:** 10.1101/2024.03.27.587031

**Authors:** Xun Zhang, Charlotte Buckley, Matthew D Lee, Margaret MacDonald, Calum Wilson, John G McCarron

## Abstract

**Background:** Endothelial cell TRPV4 channels provide a control point that is pivotal in regulating blood vessel diameter by mediating the Ca^2+^-dependent release of endothelial-derived vasoactive factors. In hypertension, TRPV4-mediated control of vascular function is disrupted but the underlying mechanisms, and precise physiological consequences remain controversial.

**Methods:** Here, using a comprehensive array of methodologies, endothelial TRPV4 channel function was examined in intact arteries from normotensive WKY and hypertensive SHR rats.

**Results:** Our results show there is a notable shift in vascular reactivity in hypertension, characterized by enhanced endothelium-dependent vasodilation at low levels of TRPV4 channel activation. However, at higher levels of TRPV4 activity, this vasodilatory response is reversed, contributing to the aberrant vascular tone observed in hypertension. The change in response, from dilation to constriction, was accompanied by a shift in intracellular Ca^2+^ signaling modalities arising from TRPV4 activity. Oscillatory TRPV4-evoked IP_3_-mediated Ca^2+^ release, which underlies dilation, decreased, while the contraction inducing sustained Ca^2+^ rise, arising from TRPV4-mediated Ca^2+^ influx, increased. Our findings also reveal that while the sensitivity of endothelial cell TRPV4 to activation was unchanged, expression of the channel is upregulated and IP_3_ receptors are downregulated in hypertension.

**Conclusions:** These data highlight the intricate interplay between endothelial TRPV4 channel expression, intracellular Ca^2+^ signaling dynamics, and vascular reactivity. Moreover, the data support a new unifying hypothesis for the vascular impairment that accompanies hypertension. Specifically, that endothelial cell TRPV4 channels play a dual role in modulating blood vessel function in hypertension.

## INTRODUCTION

Hypertension is an insidious condition linked to various vascular disorders, including coronary artery disease, stroke, dementia and renal failure. Although increased blood pressure levels are positively and continuously related to increasing cardiovascular risk, the precise mechanisms that lead from hypertension to ill health are poorly understood. Nonetheless, changes in arterial structure and function, triggered by endothelial cell dysfunction^1^, are central to hypertension’s onset and progression^2–4^.

The endothelial cell lining of blood vessels controls most cardiovascular functions. For example, the endothelium controls the regulation of inflammation, vascular remodeling^5,6^, and the moment-to-moment control of blood flow^7,8^. The endothelium regulates each vascular functionality by releasing various diffusible vasoactive molecules. These substances include anti-inflammatory and vasodilator factors (e.g., nitric oxide, prostacyclin, and endothelium-derived hyperpolarizing factor) and pro-inflammatory/vasoconstrictor factors (e.g. reactive oxygen species, prostanoids, and endothelin). Often, pathological conditions are associated with a shift from an anti-inflammatory to pro-inflammatory state, leading to vascular inflammation, narrowed blood vessels, and reduced blood flow. Whether a cause or a consequence, these alterations contribute substantially to the progression of hypertension^9,10^. However, whether the changes result from a redirection of existing signaling pathways or recruitment of new ones remains uncertain.

The release of endothelium-derived relaxing and contracting factors is triggered by changes in cytosolic Ca^2+^ concentration^11–14^. The regulation of endothelial Ca^2+^ involves both internal and extracellular sources^15^. Primarily, Ca^2+^ release from the internal store is mediated by IP_3_ receptors^16^ and several Ca^2+^ channels expressed on the plasmalemma membrane control Ca^2+^ influx. Among these influx channels, the Transient Receptor Potential Vanilloid 4 (TRPV4) is now recognized as having particular significance in regulating Ca^2+^ entry in endothelial cells^17–19^. Widely expressed in vascular endothelial cells, TRPV4 channels exhibit high Ca^2+^ permeability and respond to physical stimuli like shear stress, stretch, and intravascular pressure to promote endothelium-dependent vasodilation^14,20,21^.

In hypertension, the potential importance of TRPV4 to vascular control is increased since the physical forces that act on the endothelium (such as shear stress and intravascular pressure) are substantially altered. The modified physical forces raise the possibility that altered TRPV4-mediated Ca^2+^ influx may be a component of the endothelial changes that are associated with hypertension. In support, in a nitric oxide synthase inhibitor-induced model of hypertension, blood pressure was greater in global TRPV4 knock-out than in control normotensive mice. The authors proposed that endothelial TRPV4 channel-dependent vasodilation opposes blood pressure increases generated by the nitric oxide synthase inhibitor^22^.

Yet, despite their acknowledged importance to endothelial Ca^2+^ influx, published studies present a complicated picture of the role of endothelial TRPV4 channels in hypertension. While the physical forces that activate TRPV4 are increased, the channel’s activity is reported to be decreased in various models of hypertension^23–25^. The observations of decreased TRPV4 activity has led to the proposal that strategies to increase endothelial TRPV4 activity may restore vascular function in hypertension. In other studies, TRPV4 activity has been reported to be largely unaltered or even increased in hypertension^26^. Further confusing the situation, the alterations in TRPV4 activity that occur in hypertension may either increase vasodilation^26^, reduce vasodilation^23^ ^24^ ^25^, evoke endothelium-dependent contractions^27^ or have little effect^28^. As a result of the various reported changes in TRPV4 activity, numerous hypotheses have emerged concerning the physiological or pathophysiological implications of altered TRPV4 activity in hypertension. These include, TRPV4 may potentially serve no substantial role in the blood pressure changes^29,30^, may act as a compensatory mechanism to offset elevations in blood pressure^26^, or may be a cause that underlies the increased blood pressure that characterizes hypertension^27^.

Our study was undertaken to examine endothelial TRPV4-mediated Ca^2+^ responses in hypertension. In particular, we sought to determine whether changes in endothelial function that occur in hypertension were associated with redirection of TRPV4 responses. We show there is increased TRPV4 expression and TRPV4-mediated Ca^2+^ signaling in hypertension. In normotensive controls, TRPV4 activation induced an endothelium-dependent vasodilation at all levels of channel activity. In hypertension, TRPV4 activation generated increased vasodilator responses at low levels of channel activation. However, at higher levels of TRPV4 activation, there was reduced vasodilation generating an increased contraction. Two features explained the switch in TRPV4-evoked vasomotor responses in hypertension; 1) a reduction in IP_3_-mediated Ca^2+^ signaling and 2) increased TRPV4 Ca^2+^ influx. These results show TRPV4 channels play a dual role in hypertension. Low levels of TRPV4 activity may offer some protection while at higher levels of activity, TRPV4 may contribute to the increased vascular tone that accompanies hypertension.

## MATERIALS AND METHODS

All study data are included in the article and supporting information. An expanded Material and Methods section can be found in the Supplemental Materials.

### Animals

Animal care and experimental procedures were conducted in accordance with the relevant UK Home Office Regulations, (Schedule 1 of the Animals [Scientific Procedures] Act 1986, UK) and were approved by the University of Strathclyde Animal Welfare and Ethical Review Body. Animal studies are reported in compliance with the ARRIVE guidelines^31^.

42 Wistar-Kyoto (WKY, 7 weeks old) and 42 Spontaneously Hypertensive (SHR, 8 weeks old) rats, (purchased from Envigo, United Kingdom) were housed 3 per cage and maintained until 6 months of age at The University of Strathclyde Biological Protection Unit. All animals had ad libitum access to standard rat chow (Rat and Mouse No.1 Maintenance, 801151, Special Diet Services, UK) and water. A 12:12 light/dark cycle was used with a temperature range of 19 to 23°C (set point 21°C) and humidity levels between 45% and 65%. Animals were kept in RC2F cages (North Kent Plastic, UK) with aspen wood chew sticks and hanging huts for enrichment. At 6 months of age, animals were euthanized by cervical dislocation with secondary confirmation via decapitation in accordance with Schedule 1 of the Animals (Scientific Procedures) Act 1986. Only male rats were used to limit variability and one WKY rat died before reaching 6 months of age, and was not included in the study.

### Experimental Techniques

*In vivo b*lood pressure was monitored via tail cuff plesthysmygrophy (Visitech Systems BP-2000). Small mesenteric artery vascular reactivity and Ca^2+^ activity were studied using flat-mounted (*en face*) artery preparations or freshly isolated smooth muscle cells. For vascular reactivity experiments, we used the open source blood vessel diameter measurement software, Vasotracker Offline Analyzer^32^, and recorded vascular tone in response to various pharmacological treatments. For Ca^2+^ imaging experiments, endothelial cells were preferentially loaded with the Ca^2+^ indicator, Cal 520/AM (5 µM). Ca^2+^ responses were evoked by agonists or photolysis of caged inositol triphosphate (IP_3_). Images were acquired using various epifluorescence microscopes optimised for low-light Ca^2+^ imaging. Ca^2+^ signal analysis was performed using custom Python software^33–35^. Protein expression was visualised using immunofluorescence staining of mesenteric artery rings and the following antibodies: anti-CD31 (PECAM, # AF3628, R&D Systems, 1:1000 dilution, raised in goat); anti-TRPV4R (# ACC-034, Alomone Labs, 1:1000 dilution, raised in rabbit), anti-IP_3_R (Catalogue # 07-1210, Millipore, 1:100 dilution, raised in rabbit). Endothelial cell ion channel expression was assessed using transcriptomic analysis of single-cell RNA sequencing data from mesenteric arteries generated by Cheng et al.^36^ and Python-based bioinformatic tools.

### Statistics and data analysis

Summary data are presented in text as mean ± standard deviation (SD), and graphically as individual data points mean ± SD overlaid. Data were analyzed using independent 2-sample t-tests (with Welch’s correction as appropriate), ordinary or repeated measures two-way ANOVA with Sidak’s multiple comparisons test as appropriate and as indicated in the respective figure or table legend. All statistical tests were two-sided. A p value of <0.05 was considered statistically significant.

## RESULTS

### Enhanced TRPV4 channel activity in hypertension is a double-edged sword

We investigated the involvement of TRPV4 channels in the vascular changes occurring in a genetic model of hypertension (SHR). Notably, hypertensive SHR rats exhibited elevated blood pressure compared to normotensive WKY controls (Figure 1A). Similarly, arterial wall thickness was increased in the hypertensive compared to normotensive strain (Figure 1B).

**Figure 1.**
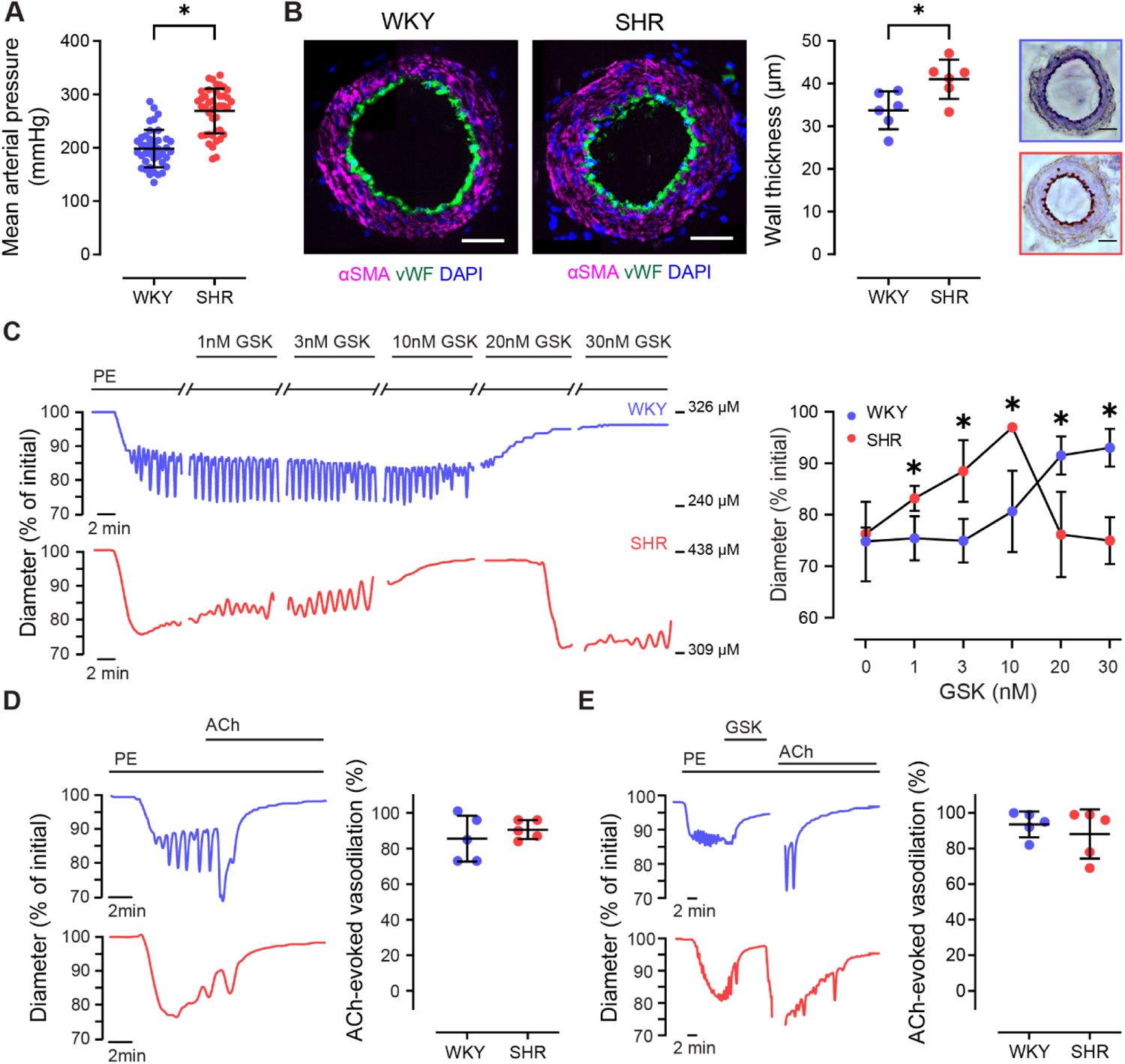
TRPV4-mediated vascular responsivity is altered in hypertension. A) Mean arterial blood pressure in spontaneously hypertensive rats (SHR) and normotensive, Wistar Kyoto (WKY), control animals. B) Representative immunofluorescence/immunohistogical images of mesenteric artery cross sections, and summary data plot showing mesenteric artery wall thickness. C-E) Representative artery diameter traces and summary data showing the effect of TRPV4 channel activation (using GSK1016790A, GSK, 30 nM) and muscarinic receptor activation (using acetylcholine, ACh, 100 nM) on vascular tone in arteries preconstricted with phenylephrine (PE, concentration adjusted to achieve ∼20% contraction; n = 5 for each). Significance markers indicate statistical significance (P < 0.05) using Student’s t-test with Welch’s correction (panels A,B,D,E) or one-way ANOVA with Tukey’s post-hoc test (C). Image scale bars = 50 µm.

To examine TRPV4-mediated regulation of blood vessel diameter in hypertension, we assessed vascular reactivity in small mesenteric arteries. In arteries from normotensive WKY, the specific TRPV4 channel activator GSK1016790A (GSK) evoked a concentration-dependent relaxation (Figure 1C). However, in the hypertensive strain, the response to GSK was biphasic. Low concentrations of GSK (≤ 10 nM) induced vasorelaxation that was significantly greater than those in the normotensive strain (Figure 1C). At higher concentrations, there was a reversal of this effect, leading to a constriction of the arteries. The TRPV4 channel blocker, ruthenium red, had no effect on PE-evoked vasoconstriction, but it effectively blocked the response to GSK (Figure S1), confirming the role of TRPV4 channels in the GSK-evoked vascular response. The observed increase in TRPV4-mediated relaxation and the biphasic response to TRPV4 activation in arteries from SHR animals (initial relaxation followed by a reversal to contraction at higher concentrations) suggests a complex regulatory mechanism involving TRPV4 in hypertension.

TRPV4-mediated vasodilation required an intact endothelium in WKY and SHR animals (Figure S2). Additionally, the reversal of relaxation at higher concentrations of GSK was not due to direct activation of smooth muscle as GSK (30 nM) did not evoke contraction in arteries in which the endothelium had been removed (Figure S3) nor evoke a Ca^2+^ increase in isolated smooth muscle cells (Figure S4A). The observed variation in TRPV4-mediated responses is specific to the TRPV4 pathway and does not reflect a general alteration in endothelial function; endothelium-dependent relaxation evoked by acetylcholine was similar in WKY and SHR rats (Figure 1D). Furthermore, vasoconstriction to higher levels of TRPV4 activation did not arise from a non-specific inhibition of endothelial cell function as endothelial reactivity to acetylcholine remained intact after the occurrence of the biphasic response (Figure 1E).

These results collectively indicate that there is increased endothelial sensitivity to TRPV4 activation in hypertension, leading to greater vasorelaxation in arteries from hypertensive animals compared to normotensive controls. Furthermore, at high levels of TRPV4 activation, arteries from hypertensive animals exhibit a secondary response that is characterized by a reversal of the initial vasorelaxation response.

### Distinct signaling pathways

Next, we sought to elucidate the mechanisms underlying TRPV4-mediated responses in hypertension. Previous findings have indicated that TRPV4 channel activity increases endothelial Ca^2+^ levels to drive vasorelaxation^18,37,38^. Given our observations that high levels of TRPV4-mediated activation reverses vasorelaxation, we speculated that endothelial Ca^2+^ concentration decreases at higher [GSK] in hypertension.

To test this hypothesis, we examined TRPV4-mediated Ca^2+^ signaling in large populations of endothelial cells using wide-field imaging (Figure 2A). Contrary to our hypothesis, TRPV4 activation generated a larger concentration-dependent increase in endothelial cell Ca^2+^ in hypertensive animals when compared to normotensive controls (Figure 2B). This surprising finding prompted the question: why do greater Ca^2+^ increases lead to a reversal of the vasorelaxation response at higher GSK concentrations in hypertensive animals but not in normotensive controls?

**Figure 2.**
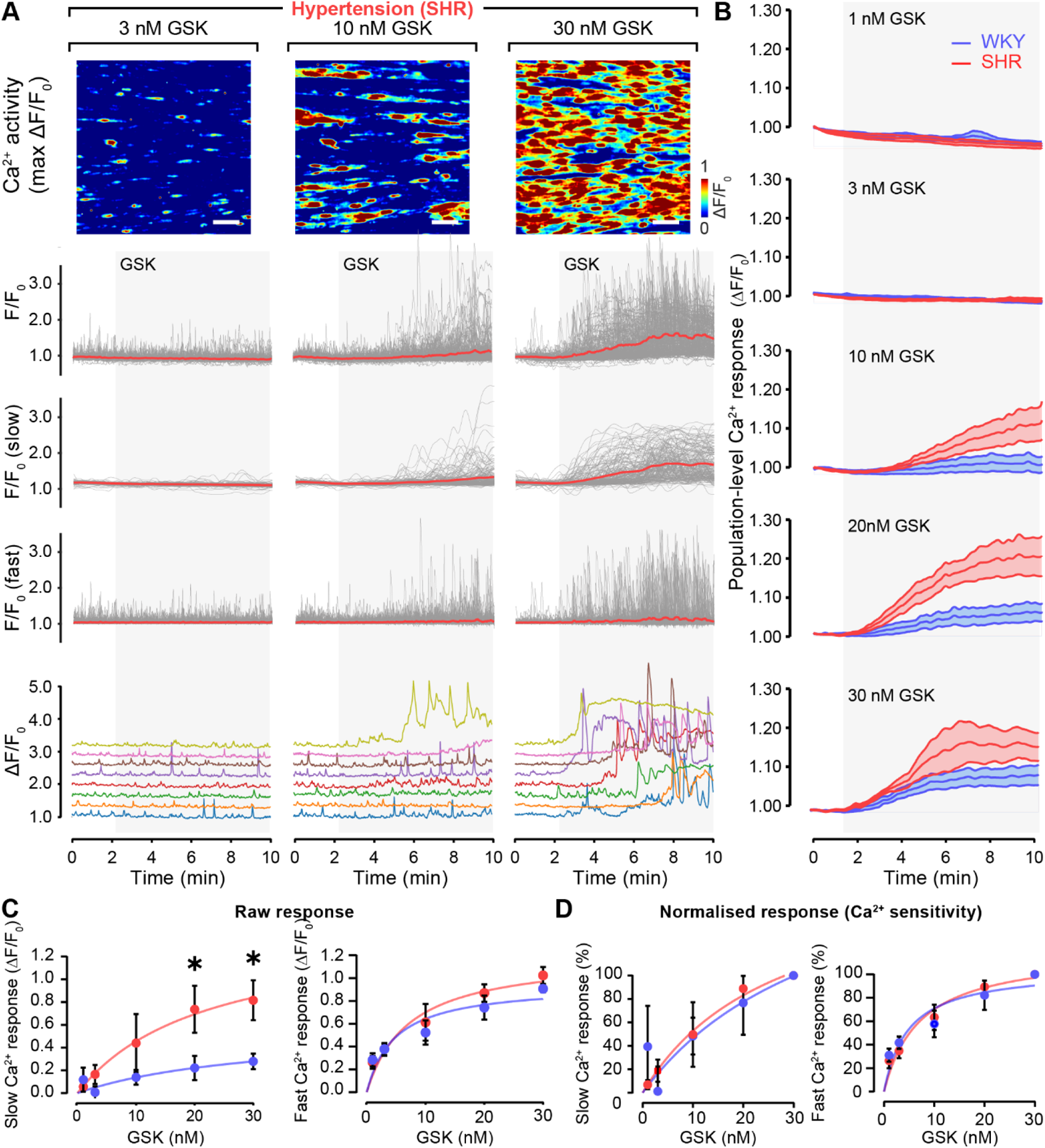
Hypertension impairs TRPV4-mediated endothelial Ca^2+^ signaling. A) Concentration-dependence of TRPV4-mediated (GSK-evoked) endothelial Ca^2+^ activity. Rows display: pseudocolored ΔF/F_0_ maximum intensity projections of a field of mesenteric endothelium (∼180 cells) stimulated with increasing concentrations of GSK; raw F/F_0_ Ca^2+^ signals and corresponding slow/fast signal component extracted from each cell; and example traces from nine separate cells. B) Mean population-level responses (F/F_0_) to increasing TRPV4 channel activation in spontaneously hypertensive rats (SHR) and normotensive, Wistar Kyoto (WKY), control animals. C–D) Summary data illustrating the concentration-dependence of the slow and fast components of the Ca^2+^ response. In D, the data has been normalized to the maximum response from each strain. Data are mean ± SD (n = 6 per group). Significance markers indicate statistical significance (P < 0.05) using two-way ANOVA with Tukey’s post-hoc test. Image scale bars = 50 µm.

The Ca^2+^ response to TRPV4 activation is composed of two components^18,37^: a slow, sustained increase in Ca^2+^ levels arising from Ca^2+^ entry through TRPV4 channels, and rapid intracellular Ca^2+^ oscillations resulting from TRPV4-mediated, IP_3_-evoked Ca^2+^ release. Given that Ca^2+^ influx is associated with endothelium-dependent vasoconstriction in hypertension^39^, whilst IP_3_-mediated Ca^2+^ activity is linked to endothelium-dependent vasodilation, we speculated that differences in the contribution of Ca^2+^ influx and Ca^2+^ release may explain our findings. Specifically, we hypothesized that the reversal of TRPV4-mediated vasodilation at high levels of TRPV4 activation occurred as a result of a shift in the dominant component of the signal moving from IP_3_-mediated Ca^2+^ increases to Ca^2+^ influx.

To investigate this hypothesis, we first isolated the influx and release components of cellular Ca^2+^ signals to examine each of their contributions to the overall response. The signals were isolated based on the kinetics of the signals; slow persistent elevation is consistent with Ca^2+^ influx while rapid oscillations with Ca^2+^ release from the internal store (Figure 2A; see Methods).

The amplitude of each component increased with the level of TRPV4 activation. Consistent with our hypothesis, the amplitude of the response was significantly higher in hypertension than in normotensive controls, whilst there was no difference in the amplitude of the fast component (Figure 2C). This alteration was not due to changes in sensitivity to the activator, GSK, as there was no difference in either component when normalized to the maximum response (Figure 2D). Once again, the results do not reflect a generalized alteration in endothelial function since endothelial Ca^2+^ responses evoked by three distinct mechanisms - muscarinic receptor activation, emptying of internal Ca^2+^ stores, and direct activation of IP_3_ receptors - were each similar in the two rat strains (Figure S4B & Figure S5).

On the basis of these observations, we next performed additional analysis on the Ca^2+^ response evoked in each cell upon TRPV4 activation (Figure 3). Our analysis revealed a distinct shift in the distribution of Ca^2+^ signaling modes in individual cells. In both WKY and SHR, the initial response to TRPV4 activation was characterized by Ca^2+^ oscillations (Figure 3A-B). However, a subset of cells transitioned from this oscillatory pattern to a sustained Ca^2+^ response with a high plateau (Figure 3C). As cells switched to a sustained response, the mean frequency of oscillations reduced as the majority of cells exhibited no discernible oscillations. These data are consistent with a transition from IP_3_-mediated Ca^2+^ release to Ca^2+^ influx. Of significant interest, the number of cells that transitioned from oscillatory responses to sustained Ca^2+^ influx was increased in hypertensive arteries (Figure 3D-E).

**Figure 3.**
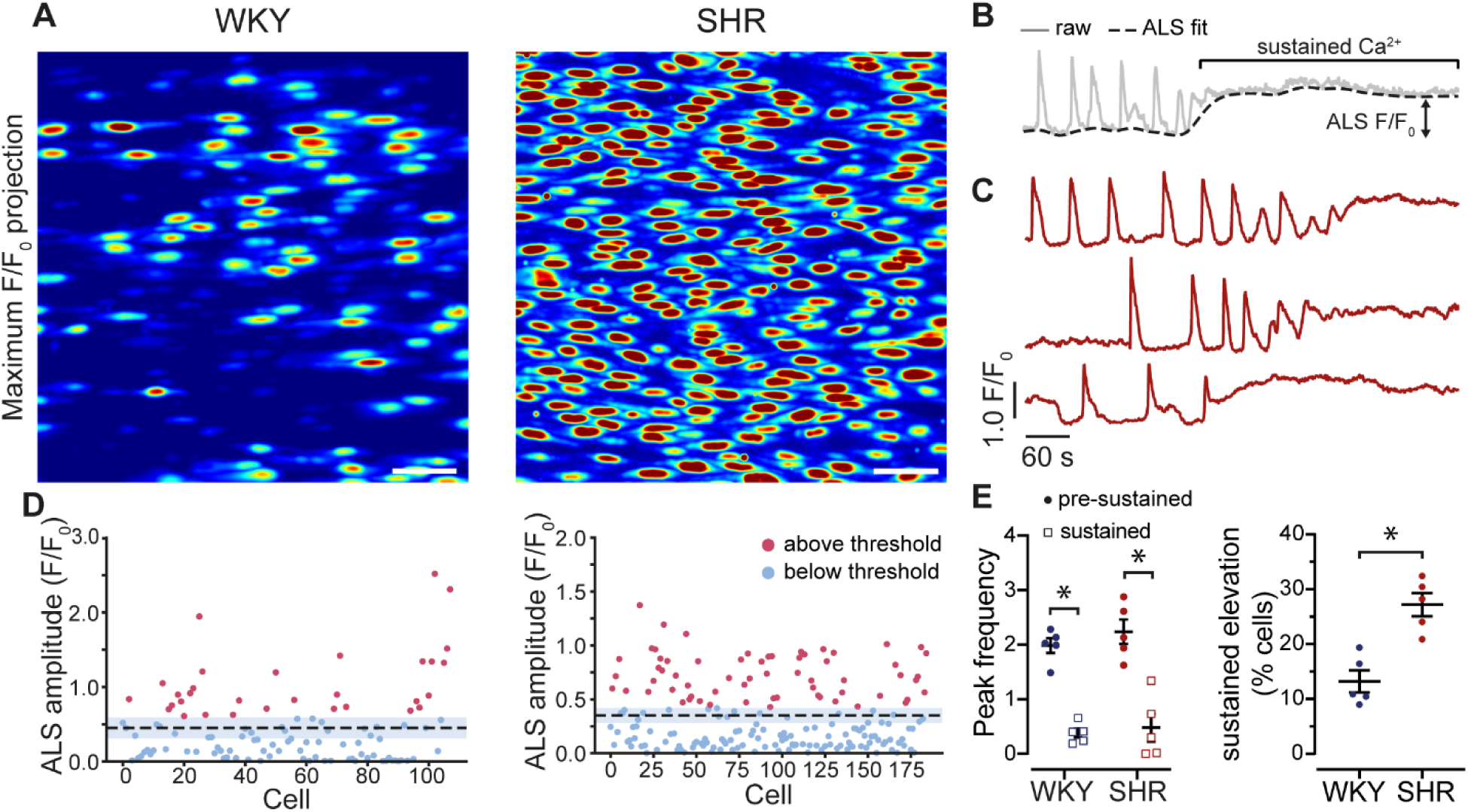
TRPV4-mediated Ca^2+^ signaling shifts from release to influx in hypertension. A) Example endothelial cell Ca^2+^ activity images (ΔF/F_0_ maximum intensity projections) illustrating TRPV4-mediated responses in hypertensive animals and normotensive controls B) Example single-cell calcium trace (grey) with asymmetric least squares (ALS) fit (black dashed line). C) individual Ca^2+^ traces from (A) that are greater than the threshold in (D), illustrating a transition from an oscillatory signal to a sustained elevation. D) Amplitude of ALS fit of all cells in (A) with mean (black dash line) and SEM (blue shade). Values above the threshold (see methods) are colored red. E) Summary data illustrating the switch from an oscillatory to a sustained elevated response to TRPV4 channel activation (left) and the percentage of cells that exhibit this switch in hypertension and normotensive controls (right). Data are mean ± SD (n = 5 per group). Significance markers indicate statistical significance (P < 0.05) using Student’s t-test with Welch’s correction. Image scale bars = 50 µm.

Collectively, our findings reveal that vasodilator responses in normotensive animals arise largely due to IP_3_-mediated Ca^2+^ activity, and that the shift to vasoconstriction in hypertension may be attributed to a transition to a sustained Ca^2+^ influx.

### TRPV4 Channel Expression is Upregulated in Hypertension

We next investigated the possibility that hypertension-induced alterations in TRPV4 function and signaling were paralleled by changes in the endothelial cell transcriptome. Specifically, we hypothesized that the expression of TRPV4 ion channels would be increased in endothelial cells of hypertensive animals. Immunostaining of endothelial cells (confirmed by the cell adhesion label CD31) revealed a diffuse expression pattern of both TRPV4 ion channels and IP_3_ receptors (Figure 4A-B). This expression pattern was not observed when the secondary antibody was omitted (Figure S6). In line with our initial hypothesis, the endothelial TRPV4 fluorescence signal appeared to be higher in hypertensive animals when compared with the normotensive control group (Figure 4A). However, IP_3_ receptor expression was reduced (Figure 4B).

**Figure 4.**
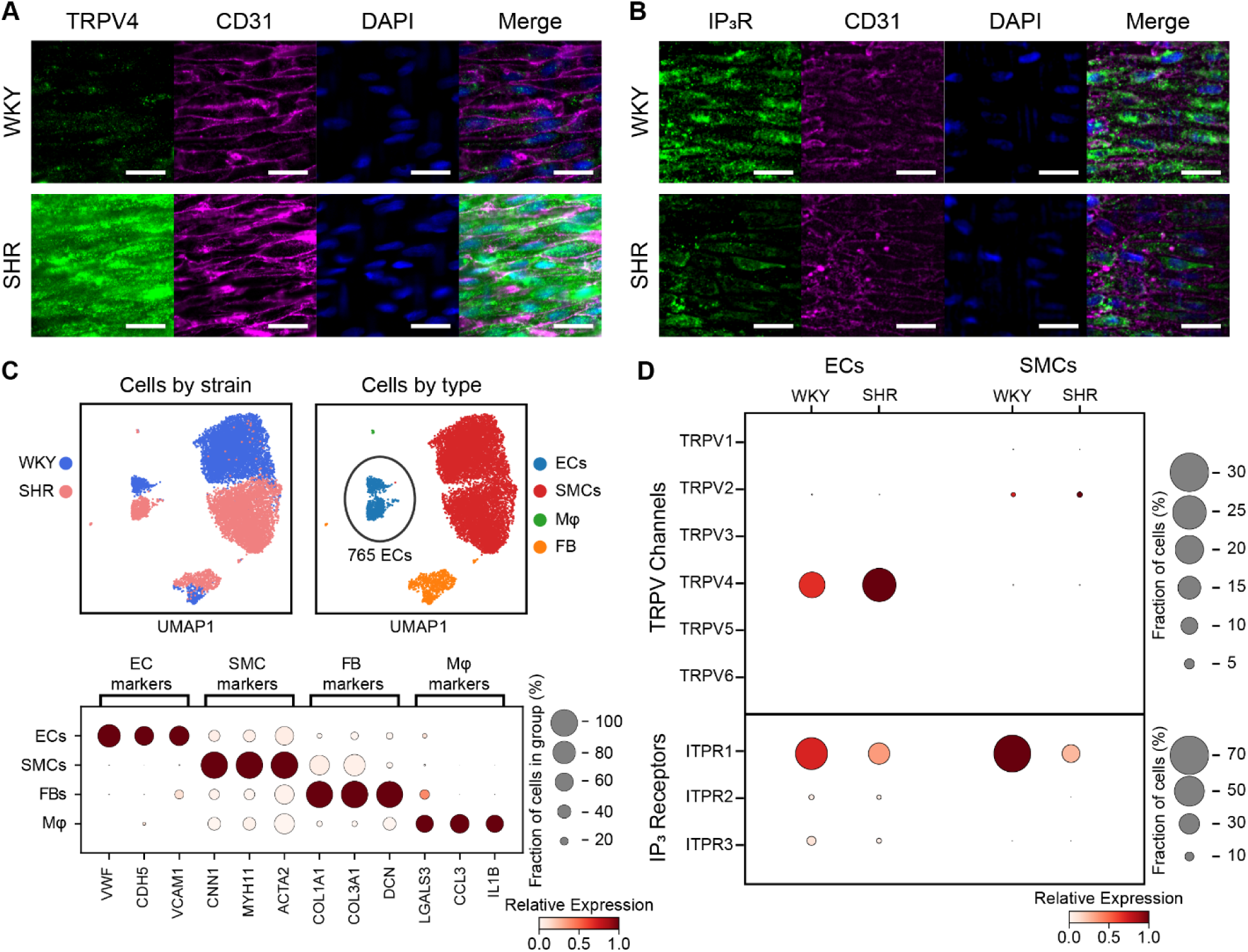
Endothelial TRPV4 channel and IP_3_ receptor expression are altered in hypertension. **A-B)** Endothelial cell ion channel expression in normotensive (WKY) and hypertensive (SHR) rats revealed by in situ fluorescence imaging. Images in show TRPV4 (panel A) or IP_3_ (panel B) receptor (green), cell-cell borders (CD31, magenta) and nuclei (DAPI, blue). No fluorescence signal was detected in arteries incubated with secondary antibodies in the absence of any primary antibody (Figure S6). Scale bars = 25 µm. C-D) Single-cell transcriptomic analysis of vascular cells from mesenteric arteries of normotensive and hypertensive rats. In total, 12,161 expression profiles were analyzed. C) Uniform manifold approximation (UMAP) projection embeddings (top) and dot plot analysis (bottom) of cell-specific marker genes. D) Dotplot of TRPV and IP_3_ receptor isoform expression in endothelial cells and smooth muscle cells. In UMAP plots, data from individual cells are colored by animal strain (WKY, blue; SHR, pink) or annotated cell type (EC, endothelial cell; SMC, smooth muscle cell; FB, fibroblasts; and Mφ, macrophages). In dotplots, the size of the dot indicates the percentage of cells in a group in which the indicated gene was detected, and the color represents the relative expression level of that gene in the indicated group. Single cell sequencing data obtained from Gene Expression Omnibus (GSE149777), see Table S1 and Table S2 for differential expression analysis^36^.

To confirm these findings, we analyzed single-cell RNA data from mesenteric arteries generated by Cheng et al.^36^. The dataset contains the expression levels of over 20,000 genes from over 12,000 mesenteric artery cells isolated from hypertensive, SHR, and normotensive, WKY rats. Using these data, we performed automated cell-type annotation and transcriptomic data from 765 high-quality endothelial cells (Figure 4C-D). These cells have robust expression of the canonical endothelial cell specific marker genes, vWF, CDH5, and VCAM1, with minimal expression of smooth muscle, fibroblast, or macrophage markers (Figure 4C). TRPV4 and IP_3_ receptor expression correlated with our immunostaining results (Figure 4D) – endothelial TRPV4 expression was ∼60% higher in hypertensive versus normotensive control rats, whilst IP_3_ receptor expression was ∼60% lower in the hypertensive strain. Additionally, TRPV4 channel expression was not detected in smooth muscle cells.

Together, these results suggest that the augmented vascular responses observed in hypertension arise from upregulated expression of plasmalemmal TRPV4 channels. The decreased expression of IP_3_ receptors may explain the reduction in endothelial control of basal vascular tone previously reported to occur in hypertensive rats^40^.

## DISCUSSION

Here we show there is increased TRPV4 channel expression and activity in hypertensive animals when compared to normotensive controls. Since there is no change in TRPV4 sensitivity to pharmacological activation, increased expression of the channel appears sufficient in explaining the heightened Ca^2+^ signaling on channel activation. The consequences of increased TRPV4-mediated Ca^2+^ signaling in hypertension are complex. At low levels of channel activation, there is increased vasodilation when compared to normotensive controls. This observation is consistent with the increased expression of the channel and Ca^2+^ signaling that occurs in hypertension. However, paradoxically, the increased Ca^2+^ signaling that occurs at higher levels of channel activation resulted in a reversal of endothelium-dependent dilation and vascular contraction. These results suggest that altered TRPV4 activity in hypertension may either be protective, at low levels of channel activity, or a contributor to the progression of hypertension, at high levels of channel activity.

The question arises as to why different types of contractile response occur to the Ca^2+^ signaling events evoked by TRPV4 activation in hypertension? An explanation is found in the distinctive nature of the Ca^2+^ signals generated at low and high levels of TRPV4 activity. The signals generated by TRPV4 activation consist of two major components: a slow sustained response and rapid transient Ca^2+^ oscillations. The slow component arises from Ca^2+^ influx via TRPV4 which generates the sustained Ca^2+^ increase. At low levels of activation this sustained rise is relatively small. The sustained rise is amplified by Ca^2+^-induced Ca^2+^ release acting at IP_3_ receptors. It is the latter that generates the rapid transient Ca^2+^ oscillations^18,41^. These oscillations occur at all levels of TRPV4 activation in normotensive controls. In hypertension, the oscillations have prominence at lower levels of activation, whereas the sustained slow component dominates at higher levels of TRPV4 activity. It is tempting to speculate, therefore, that dilation is associated with rapid Ca^2+^ oscillations while contraction is promoted by the increased sustained response. Thus, there appears to be a redirection of Ca^2+^ signals to generate a different functional outcome (contraction) at higher levels of channel activity in hypertension.

The altered priority of the slow and rapid types of Ca^2+^ signal in hypertension can itself be explained by changes in the expression of TRPV4 and IP_3_ receptors. In hypertension there is increased expression of TRPV4 and, significantly, a decreased expression of IP_3_ receptors. It is IP_3_ receptors that underlies the Ca^2+^ oscillations and dilation. These results suggest that, in hypertension, as the extent of TRPV4 activation increased, so does the slow sustained component. At the same time as the sustained component increases, the relative contribution from rapid oscillations associated with IP_3_ receptors decreases because of the reduced expression of the receptor.

In hypertension, at lower levels of TRPV4 activity, there is increased dilation when compared to normotensive controls. The IP_3_ response, which underlies the relaxation, is increased. Yet IP_3_ receptor expression is reduced, and the store content is unaltered in hypertension. The event that triggers IP_3_-evoked Ca^2+^ release is Ca^2+^ influx via TRPV4. TRPV4 expression is substantially increased in hypertension. It seems likely that the increased event triggering IP_3_-evoked Ca^2+^ release (TRPV4 mediated Ca^2+^ influx) is sufficient to offset the decreased expression of IP_3_ receptors – at least at lower levels of activation.

Since TRPV4 channels are a major route for Ca^2+^ entry in endothelial cells they are widely regarded as promoters of nitric oxide production and in activating small- and intermediate-conductance Ca^2+^-sensitive potassium (IK and SK) channel activity. Generation of nitric oxide and activation of IK and SK results in vasodilation and decreased vascular resistance. In hypertension, there is chronically-decreased nitric oxide production and reduced SK and IK channel activity which leads to impaired endothelial dependent dilation. These observations resulted in the hypothesis that decreased Ca^2+^ entry via TRPV4 may be a component of the changes associated with hypertension. Yet the relationship between Ca^2+^ entry via TRPV4 channels and the vascular changes that accompany hypertension are disputed. There are reports of changes in the channel’s activity that result in increased vasodilation, decreased vasodilation, increased contraction or no contribution at all. For example, in hypertension, TRPV4 expression may be reduced and, as a result, channel activity impaired. The decreased channel activity is proposed to contribute to the increased vascular resistance that leads to elevated blood pressure since channel activity was associated with vasodilation^23–25^. However, in animals in which the channel has been knocked-out, blood pressure may, paradoxically, be lower than controls^28^. In other studies, rather than being decreased, TRPV4 expression and channel activity may be increased in hypertension. The increased TRPV4 activity promotes vasodilation and is proposed to provide protection against the vascular changes that occur in hypertension^26^. Alternatively, rather than underlying dilation, increased TRPV4 expression and Ca^2+^ influx may promote endothelium dependent contraction in hypertension. The increased activity of the channel and its associated contraction is proposed to be a contributory factor that leads to the increased vascular resistance that characterizes hypertension^27^. In yet other studies, TRPV4 channel knockout mice do not show increased blood pressure under resting conditions as may be expected if the channel is significant in regulating vascular resistance^29,30^. While many studies show a parallel association between altered expression of TRPV4 and functional outcomes, some studies report unaltered expression (as measured by current density) but diminished functional outcome as a result of decoupling of the TRPV4 from an effector (the KCa2.3 channel) in hypertension^42^.

Thus, there is an uncertain relationship between TRVP4 activity and the vascular changes that occur in hypertension. Our study offers at least a partial explanation for the apparently contradictory findings by highlighting various ways in which TRPV4 may alter arterial activity in precisely the same artery. We show that oscillatory Ca^2+^ signals associated with IP_3_-evoked Ca^2+^ release, triggered by TRPV4 activation, generate relaxation. The sustained Ca^2+^ signals occurring at high levels of TRPV4 activation in hypertension appear to elicit contraction as a result of reduced endothelium-dependent relaxation. It is the increased expression of TRPV4 and decreased expression of IP_3_ receptors that predisposes the artery to sustained Ca^2+^ rises and contraction that occurs at high levels of TRPV4 activation in hypertension.

The question arises as to how TRPV4-mediated Ca^2+^ entry into the endothelial cytoplasm can trigger arterial relaxation in some conditions and decreased relaxation and contraction in others? Ca^2+^ has the potential to activate numerous, and at times, conflicting processes in cells. Ca^2+^-dependent functional activities are normally coordinated to avoid conflict via timely and spatially-variant Ca^2+^ signals that are matched to control observable responses. For example, the extent of activation of effectors may rely on highly local Ca^2+^ elevations, or on the duration or amplitude or repetitiveness of transient global or local increases in the concentration of the ion. The specific outcomes resulting from increased Ca^2+^, such as effects on calmodulin, nitric oxide synthase, phospholipase A2 activation, and calmodulin-dependent kinase II, depend on the rates of ion binding to the effector (including on and off rates). The temporal aspects of Ca^2+^ concentration change, coupled with effector on and off rates, play a crucial role in determining whether or not the timeframe permits accumulated activity to encode both frequency and amplitude, as shown for NFAT translocation^43^. The large number of Ca^2+^-binding proteins with unique on and off rates for Ca^2+^ binding is critical in determining functional outcomes as appears to be the case in the present study.

Ca^2+^ activated increases in nitric oxide production and SK and IK channel activity trigger relaxation. An endothelium-dependent contraction, or decreased endothelium-dependent relaxation, may be evoked by several Ca^2+^-dependent mechanisms. Sustained Ca^2+^ rises, for example, may result in significant mitochondrial Ca^2+^ uptake. Excessive mitochondrial Ca^2+^ uptake leads to an increase in reactive oxygen species (ROS) production and oxidative stress. ROS and oxidative stress may limit relaxation by decreasing the concentration of nitric oxide through the formation of the peroxynitrite anion. The production of additional endothelium-derived contracting factors, such as prostanoids, and endothelin are also Ca^2+^ dependent. Rises in Ca^2+^ increase expression of preproendothelin-1 mRNA *via* a Ca^2+^/calmodulin/calmodulin kinase pathway ^44,45^ or prostanoids via Ca^2+^-dependent phospholipase A_2_ activation^46^.

In the present study, it seems likely that TRPV4 suppresses relaxation at higher levels of activation in hypertension (rather than cause contraction) because TRPV4 activation did not cause contraction by itself in an artery that was not pre-constricted (Figure S3). Furthermore, all vascular effects of TRPV4 in the present study are mediated via the endothelium since channel activation evoked no change in arterial tone (either contraction or relaxation) in the absence of the endothelium.

### Perspectives

The proposed importance of TRPV4 to endothelial Ca^2+^ entry and changes in vascular disease, have resulted in the channel being linked to the vascular changes in hypertension. However, in various studies the changes in TRPV4 activity has been reported as being both protective, or a contributor to the pathological changes that occur in hypertension. Our results help reconcile the contradictory proposals by highlighting two responses to TRPV4 activation in hypertension. At low levels of TRPV4 activity, there is increased relaxation to the channel’s activation in vessels from hypertensive animals when compared to normotensive controls. However, at higher levels of TRPV4 activation, endothelium-dependent dilation was impaired resulting in increased contraction. These results show that at low levels of activity, TRPV4 activity may offer some protection to the vascular changes in hypertension while at higher levels of activity, TRPV4 may contribute to the increased vascular tone that accompanies hypertension. High levels of TRPV4 activity may occur in hypertension, as a result of the increased mechanical stimuli generated by increased pressure and flow velocity in small vessels, which may contribute to the vascular changes underlying hypertension.

## Supplementary Materials

### Detailed Materials and Methods

#### Blood Pressure Measurement

Blood pressure was monitored using tail cuff plesthysmygrophy (Visitech Systems BP-2000). In each measurement session, ten measurements of heart rate, and systolic, diastolic and mean blood pressures were taken and the average of each determined. A total of 4 blood pressure determinations were made. In the initial measurements a minimum of 48hrs was left between recordings. Two further sets of blood pressure measurements were taken subsequently, one of which was the week before euthanasia. Animals were euthanized at 6 months old (∼350g) by cervical dislocation.

#### Chemicals

The physiological saline solution (PSS) used in all functional experiments consisted of:-145 mM NaCl, 2 mM MOPS, 4.7 mM KCl, 1.2 mM NaH2PO4, 5 mM Glucose, 0.02 mM EDTA, 1.17 mM MgCl, 2 mM CaCl, pH 7.4. All reagents included in the PSS were obtained from Sigma.

Cal-520/AM was obtained from Abcam (UK). Caged-IP_3_ (caged-IP_3_ 4,5-dimethoxy-2-nitrobenzyl) and Pluronic F-127 was obtained from Sichem (Germany). GSK1016790A, acetylcholine, ionomycin, cyclopiazonic acid, phenylephrine, sodium nitroprusside, ruthenium red, caffeine, and dimethyl sulfoxide (DMSO) were obtained from Sigma Aldrich (USA). All solutions were prepared fresh each day and chemicals were diluted to the desired concentration with PSS.

#### En face artery preparation

Either second or third order mesenteric arteries were used for measurement of vascular reactivity or endothelial cell Ca^2+^ signaling. Immediately following euthanasia, the mesentery bed was removed and placed in PSS. Arteries were dissected rapidly, cleaned of connective tissue and fat and used immediately. Arteries were pinned to either (1) the Sylgard-coated base of a custom chamber for use on an upright microscope or (2) a Sylgard block that was subsequently placed in a custom chamber for use on an inverted microscope. The arteries were then cut open longitudinally and pinned flat with the endothelial layer facing upwards. Endothelial cells were preferentially loaded with the fluorescent Ca^2+^ indicator Cal-520/AM (5 µM with 0.04% Pluronic F127 and 0.26% dimethyl sulfoxide [DMSO] in PSS) at 37°C for 30 minutes. In a subset of experiments, endothelial cells were also loaded with a membrane-permeant, photolabile form of IP_3_. In these experiments, cIP_3_ was included in the Ca^2+^-indicator solution. Following a 30-minute incubation period, arteries were gently washed PSS and then positioned on a microscope for imaging.

#### Smooth Muscle Cell Isolation

First to fourth order mesenteric artery segments were enzymatically digested to obtain freshly isolated smooth muscle cells. In brief, vessels cut open and then into small strips of approximately 2 mm length, and then enzymatically digested using collagenase (Type 2, 256 units/mg, 2 mg.ml^-1^) in a water bath at 37°C for 45-60 mins. The supernatant was then removed gently and the arteries triturated using a wide-bored, fire-polished glass pipette. Cells were transferred to a glass-bottomed chamber for Ca^2+^ imaging, stained with Cal-520/AM (5 µM, 30 mins, 37°C) for Ca^2+^ imaging experiments.

#### Immunocytochemistry

Arteries and mounted into Sylgard-lined 6-well plates as *en face* arterial preparations. The arteries were fixed in 4% paraformaldehyde (PFA; Agar Scientific, UK) in phosphate buffered saline (PBS) for 20 mins at room temperature. Preparations were then washed three times in glycine solution (0.1 M), three times in PBS and then permeabilized with Triton-X100 (0.2% in PBS) for 30 minutes. Cells were again washed three times in PBS, three times in antibody wash solution (150 mM NaCl, 15 mM Na_3_C_6_H_5_O_7_, 0.05% Triton-X100 in milliQ water), and incubated for one hour with blocking solution (5% donkey serum in antibody wash solution) at room temperature. All individual wash steps were 5 minutes in duration. Preparations were then incubated overnight at 4°C with combinations of goat anti-CD31 (CD31/PECAM; R&D Systems cat. #AF3628, 1:1000, raised in goat) and anti-TRPV4R (Alomone Labs, Cat. # ACC-034, 1:1000, raised in rabbit) or anti-IP_3_R (Millipore, Cat. # 07-1210, 1:100, raised in rabbit) primary antibodies diluted in antibody buffer (1:1000 dilution; 0.15 M NaCl, 15 mM Na_3_C_6_H_5_O_7_, 2% donkey serum, 1% BSA, 0.05% Triton X-100 in milliQ water). Following primary antibody incubation, preparations were washed three times in antibody wash solution, and incubated for one hour at room temperature with fluorescent secondary antibodies conjugated to Alexa Fluor 488 (donkey anti-goat, 1:1000; A-11055 labelling anti-CD31) and Alexa Fluor 555 (donkey anti-rabbit, 1:1000; A-31572 labelling anti-TRPV4) in antibody buffer. The preparations were then washed three times in antibody wash solution, incubated with the nuclear stain, 4′,6-diamidino-2-phenylindole (DAPI; 4 nM) for 5 mins, and finally washed three more times in PBS (5 mins) prior to imaging. All samples were processed in a single immunostaining run and were imaged using the same microscope settings.

#### Assessment of vascular reactivity

Vascular reactivity was assessed in isolated mesenteric arteries mounted *en face*^38,40^. Arteries were visualized at 5 Hz using an upright fluorescence microscope (FN-1; Nikon, Japan) equipped with a 16X, 0.8 numerical aperture objective, 460 nm LED illumination (CoolLED, UK), and an iXon 888 (Andor, UK) electron multiplying CCD camera. The resulting 832 x 832 µm field of view allowed quantification of vascular reactivity in opened arteries using VasoTracker edge-detection algorithms^32^. Contraction data are represented as the percent reduction from resting diameter. Relaxation data (from constricted diameter) are represented as the percent of maximal relaxation (constricted diameter to resting diameter).

Arteries were partially constricted with phenylephrine added to the perfusate (to ∼80% of resting diameter i.e. 20% contraction; ∼2 µM phenylephrine). This level of constriction enables the vessels to either dilate or constrict further under experimental pharmacological studies^18^. Arteries were then assessed for endothelium-dependent relaxation to the muscarinic receptor agonist, acetylcholine (100 nM). All arteries exhibited a relaxation greater than 70% of the maximum possible, were considered viable and used in subsequent experiments. Following washout, arteries were preconstricted once more and then responses to increasing concentrations of the TRPV4 channel agonist, GSK1016790A (GSK), were examined. In a subset of experiments, the effect of endothelium removal was assessed. In these experiments, the endothelium was removed by gently scraping the intimal surface with a fine hair and smooth muscle cell viability was confirmed using phenylephrine and sodium nitroprusside (100 µM). In an additional series of experiments, the effect of the TRPV4-channel blocker, ruthenium red (5 µM), on GSK-evoked vascular responses was assessed. In these experiments, arteries were first preconstricted with phenylephrine. Ruthenium red was then added to the perfusate for 10 minutes prior to the addition of GSK1016709A. All arteries used for experimentation had a luminal diameter ∼150 µm, and were perfused with PSS (37°C) at a rate of 1.5 ml min^-1^ using a Gilson Minipuls 3 peristaltic pump.

#### Endothelial cell Ca^2+^ imaging

Intact mesenteric artery endothelial cell Ca^2+^ activity was recorded at 10 Hz using one of three imaging systems optimized for high-resolution Ca^2+^ imaging: an upright epifluorescence microscope (described above; FN-1, Nikon, Japan) equipped with a 16X 0.8 NA objective lens, and two inverted fluorescence microscopes (TE300 or Ti-Eclipse, Nikon, Japan) each equipped with 40X and 100X oil-immersion (1.3 NA; S Fluor) objective. The TE300 microscope was also equipped with a flash lamp for localized spot photolysis (00-325-JML-C2; Rapp Optoelectronics, Germany). All microscopes were fitted with multi-band LED illumination systems (pE-300 or pE-4000; CoolLED, UK), multi-band filter cubes (UV/FITC/TRITC), and sensitive, large-format EMCCD camera (iXon Ultras, 1024 by 1024 pixels; Andor, UK). All images were acquired using µManager microscope control software^47^.

#### Ca^2+^ imaging protocols

In all experiments, responses to the muscarinic receptor agonist, acetylcholine (100 nM, perfused at 1.5 ml min^-1^), were first assessed to confirm endothelial viability and physiological function. Following washout of acetylcholine, and a 10-minute re-equilibration period, endothelial cell Ca^2+^ response to increasing concentrations of GSK were then examined. In experiments assessing Ca^2+^ store content, arteries were perfused with Ca^2+^ free PSS for 5 min to remove external Ca^2+^. Cyclopiazonic acid (20 µM) was then added to the Ca^2+^-free perfusate. In experiments examining Ca^2+^ responses evoked by direct activation of IP_3_ receptors, the inositide was released (caged IP_3_) using a UV flash of ∼ 1 ms duration^48,49^. A broadband light source coupled to the to the epi-illuminator allowed the position of the uncaging region (∼ 70 µm diameter) and which endothelial cells were directly activated by the spot photolysis system to be determined. Three recordings of responses to uncaging of IP_3_ were recorded from each biological replicate, with a 15 mins recovery period between flashes to allow Ca^2+^ store refilling to occur.

In all Ca^2+^ imaging experiments, endothelial cells were activated after a minimum baseline recording of 1 minute in duration (pharmacological activation) or 30 seconds (spot photolysis). All images were acquired at 10 Hz.

#### Analysis of Ca^2+^ activity

Single-cell endothelial Ca^2+^ activity was assessed as previously described^18,50,51^. In brief, we used automated algorithms to extract fluorescence intensity as a function of time from circular regions of interest (∼15 µm diameter) centered on each cell in our images. Fluorescence signals were then smoothed using a Savitzsky-Golay (21 point, third-order) filter, and expressed as fractional changes in fluorescence (F/F_0_) from baseline (F_0_). The baseline was automatically determined by averaging the fluorescence intensity of the 100-frame portion of each trace that exhibited the least noise. We then calculated the discrete derivative (d(F/F_0_)/dt) of each Ca^2+^ signal, and used a peak-detection algorithm to identify increases in fluorescence intensity that rose at least 15 SD above baseline noise. For acetylcholine-evoked responses, Ca^2+^ activity was quantified using the number (or percentage) of cells exhibiting spiking activity, and the amplitude of these Ca^2+^ spikes (ΔF/F_0_). Analysis of IP_3_-evoked Ca^2+^ activity was restricted to those cells in which cIP_3_ was released by applying a mask restricted to the photolysis region.

The response to TRPV4 channel activation involves two main components: (1) a slow, sustained elevation in baseline Ca^2+^ levels and (2) rapid intracellular Ca^2+^ waves (oscillations). To separate these components, each signal’s slow persistent elevation was isolated using an asymmetric least squares (ALS) smoothing technique. The fast-oscillatory signal component was separated by normalizing each signal with its ALS-smoothed counterpart to eliminate slow drifts, and signals processed as above. To give an indication of the percentage of cells exhibiting sustained elevations in Ca^2+^ levels, we calculated the start of a sustained elevation in Ca^2+^ as 90% of the ALS slope. The mean amplitude and frequency of events were measured from this point until the end of the recording. The mean amplitude (+3x standard error of the mean) was used as a threshold to define cells with the highest sustained response.

To assess the internal store content (experiments using cycolopiazonic acid), entire field-of-view average Ca^2+^ signals were extracted using ImageJ^52^ and plotted in Origin Pro (OriginLab Corporation, US). In brief, Ca^2+^ traces were plotted and the area under the curve (AUC) calculated as a measure of total internal store content.

For graphical representations of endothelial cell Ca^2+^ activity, we created single image representations of Ca^2+^ recordings. This image was created by taking the maximum intensity of F/F_0_ image sequences for the duration of the recordings. In the case of caged IP_3_-evoked Ca^2+^ experiments, the image was created from the first 2 s (20 frames) immediately following IP_3_ uncaging and presented using a JET LUT.

#### Single cell RNA Sequencing Data Analysis

We obtained Smart-Seq2 RNA sequencing libraries of single mesenteric artery cells submitted by Cheng *et al.* from the Gene Expression Omnibus database (GSE149777)^36^. The dataset contains the read count matrix of 25340 genes from cells isolated from arteries from WKY (7197 cells) and SHR (6549 cells) rats (n = 7-8 pooled animals per group). All analysis was performed using custom Python scripts. First, data were merged into a single AnnData object with disease status/strain annotation using the Scanpy package^53^. To ensure the inclusion of only high-quality cells for downstream analysis, cells with fewer than 2100 Unique Molecular Identifiers or expressing less than 500 or more than 3500 genes were excluded to remove low-quality or potential doublet cells, and those with over 15% mitochondrial gene expression were also discarded. Raw counts were converted to log counts per 10,000 by log-normalization and subsequently scaled. To focus on the most informative features, we identified highly variable genes based on specified mean expression and dispersion thresholds (minimum mean of 0.0125, maximum mean of 3, and minimum dispersion of 0.5). After filtering, we corrected for confounding factors by regressing out effects attributable to total counts and mitochondrial gene expression percentages, scaled the data, performed Principal Component Analysis (PCA) for dimensionality reduction, and embed the cells in a two-dimensional space using the Uniform Manifold Approximation and Projection (UMAP) technique.

Cell clusters were identified using the Leiden algorithm (resolution parameter = 0.5). To identify specific cell types, we used Over-Representation Analysis against the PanlaoDB markers for canonical cell types. The technique identified a total of 12171 cells (10545 smooth muscle cells, 844 fibroblasts, 766 endothelial cells, and 16 macrophages). To verify the cell-type annotations, we constructed dot plots for selected marker genes of endothelial cells (vWF, CDH5, VCAM1) smooth muscle cells (CNN1, MYH11, ACTA2), fibroblasts (COL1A1, COL3A1, DCN), and macrophages (LGALS3, CCL3, IL1B). To assess for alterations in the expression of specific ion channels attractable to genetic variances, we subset the data and performed differential expression analysis using a t-test, grouped by animal strain.

### SUPPLEMENTARY FIGURES

**Figure S1.**
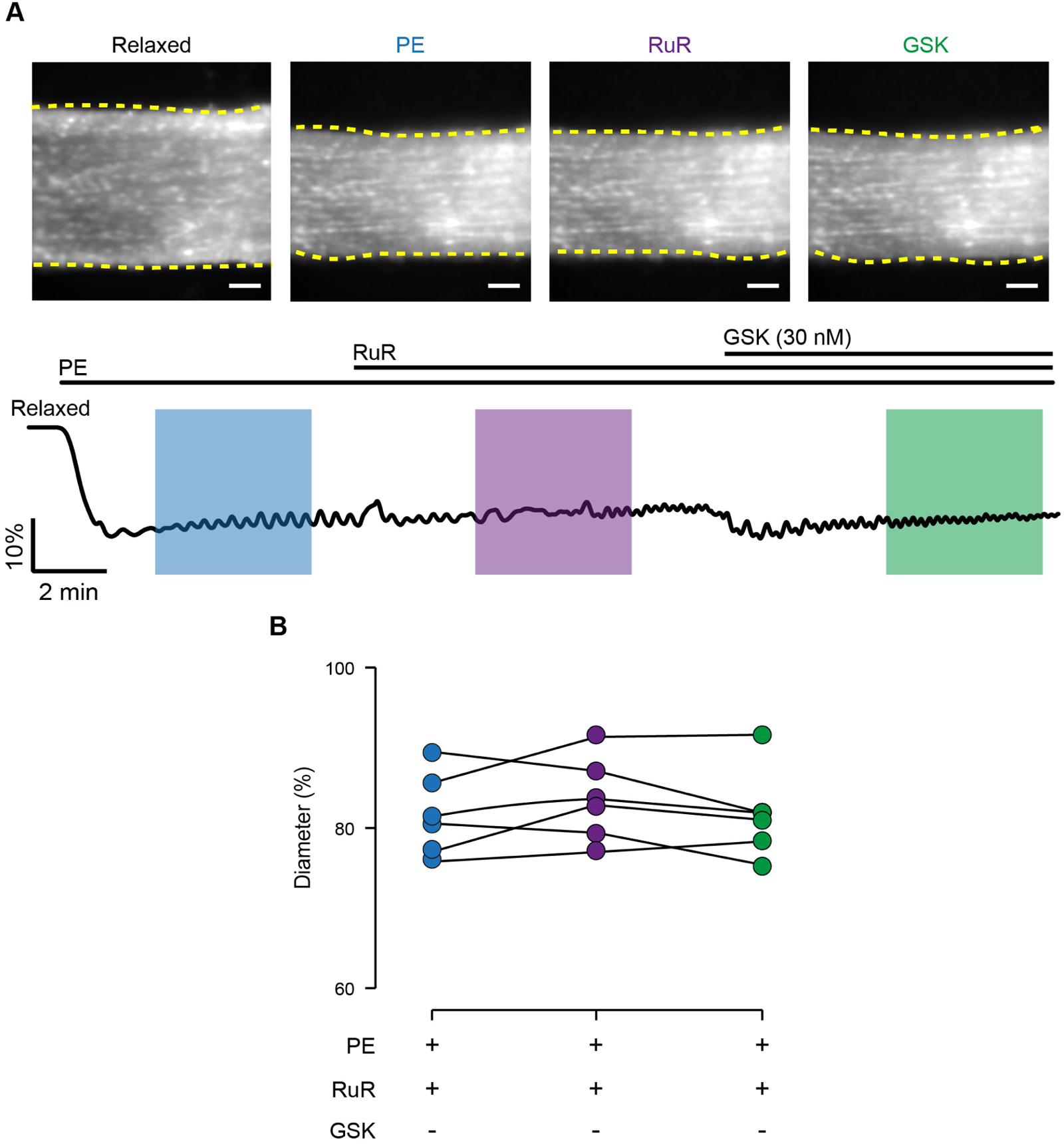
TRPV4 inhibition does not reverse PE-induced contraction. A-B) Still frame images/time course (A) and summary data (B) showing the effect of the TRP channel inhibitor, ruthenium red (5 μM), and the selective TRPV4 channel agonist, GSK1016790A (GSK, 30 nM), on mesenteric artery tone during vasoconstriction evoked by phenylephrine (PE, 2 μM). Data were assessed using a one-way ANOVA for paired data with Tukey’s post-hoc test for multiple comparisons. Image scale bars = 100 µm.

**Figure S2.**
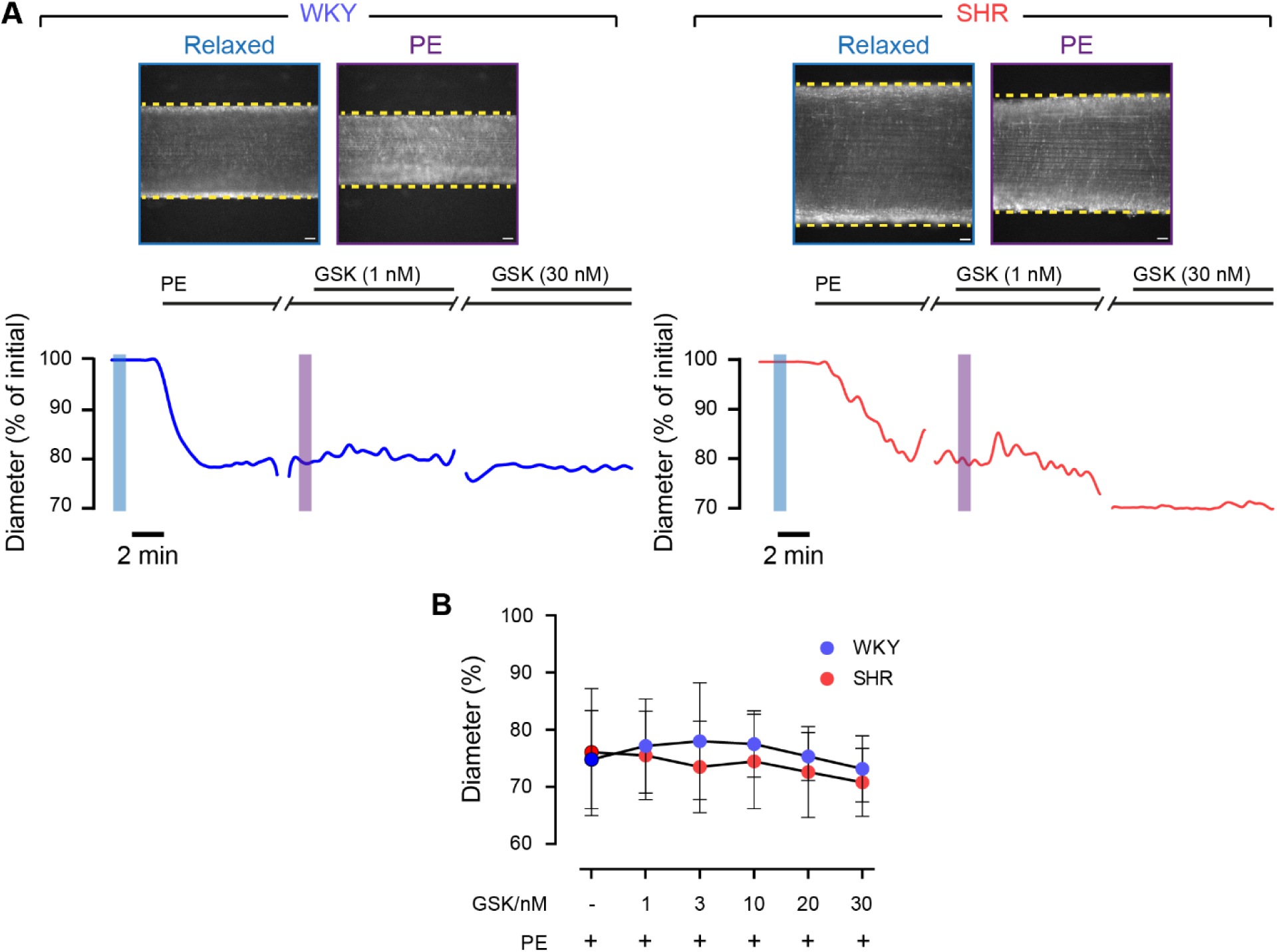
TRPV4-mediated regulation of vascular tone is endothelial dependent. A-B) Still frame images/time course (A) and summary data (B) showing the effect of the TRPV4 channel agonist, GSK1016790A (GSK), on mesenteric artery tone during vasoconstriction evoked by phenylephrine (PE, 2 μM). Endothelial cells were mechanically removed with a fine hair, and these arteries remained capable of constricting to phenylephrine (PE) and relaxing to the nitric oxide donor, sodium nitroprusside (SNP; see Figure S3). Image scale bars = 50 µm.

**Figure S3.**
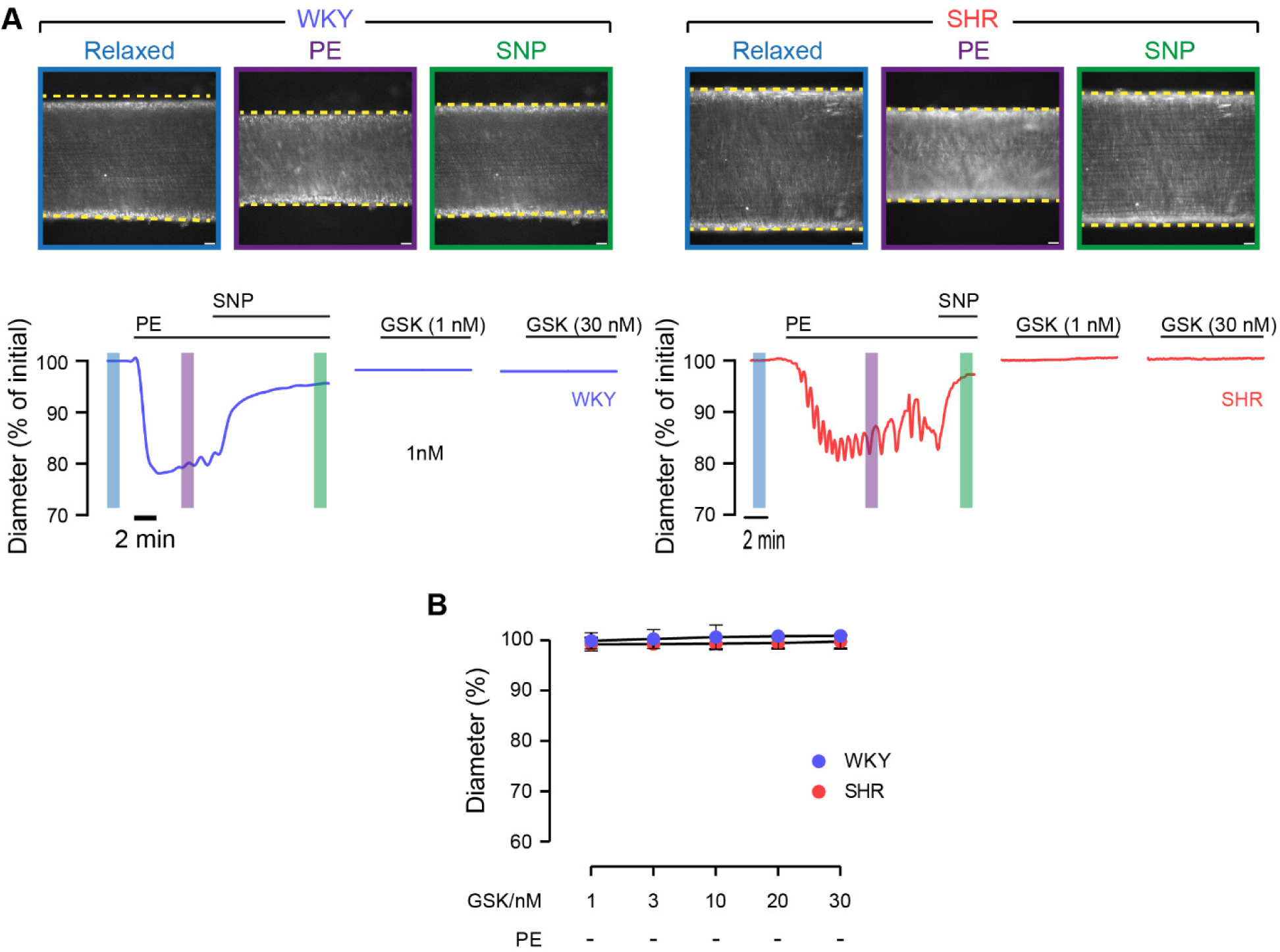
TRPV4-mediated vasoconstriction is absent in endothelium-denuded arteries. A-B) Still frame images/time course (A) and summary data (B) showing the effect of the TRPV4 channel agonist, GSK1016790A (GSK), on the resting tone of mesenteric arteries which lacked a functional endothelial cell layer. Endothelial cells were mechanically removed with a fine hair, and these arteries remained capable of constricting to phenylephrine (PE) and relaxing to the nitric oxide donor, sodium nitroprusside (SNP). Image scale bars = 50 µm.

**Figure S4.**
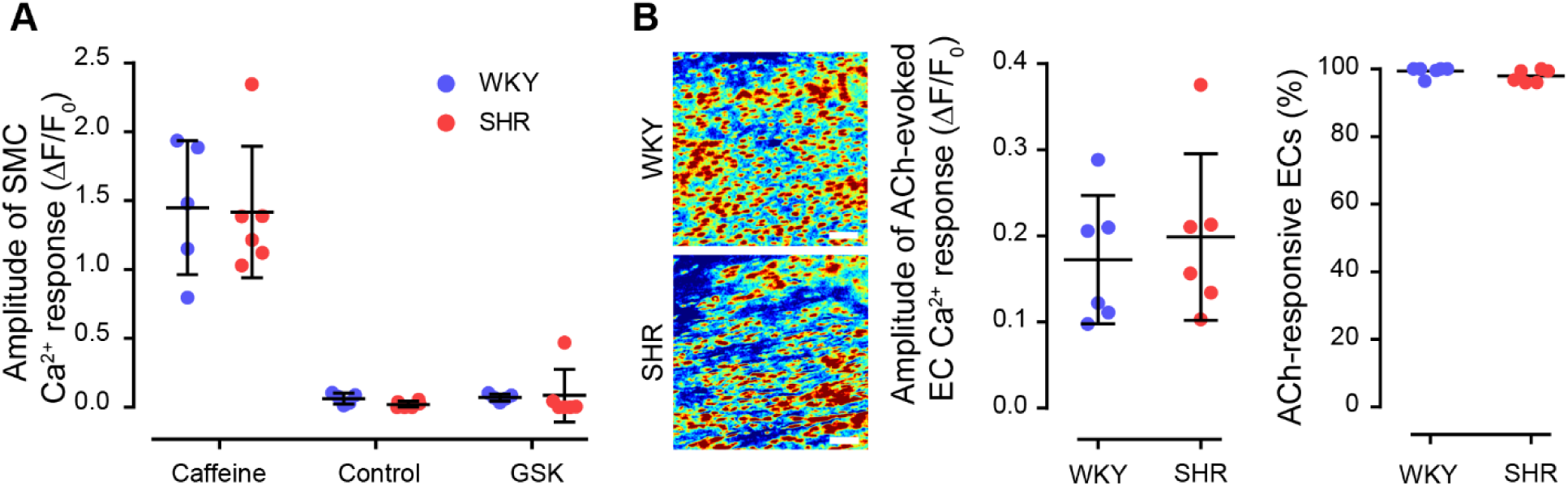
TRPV4 is not functional in isolated smooth muscle cells and acetylcholine-evoked responses are maintained in hypertension. A) Summary data showing the amplitude of agonist-evoked Ca^2+^ responses in isolated mesenteric artery smooth muscle cells from normotensive (Wistar Kyoto, WKY) and hypertensive (spontaneously hypertensive rat, SHR) animals. B) Example endothelial cell Ca^2+^ activity images (ΔF/F_0_ maximum intensity projections) and summary data illustrating ACh-evoked Ca^2+^ activity in the hypertensive animals and normotensive controls.

**Figure S5.**
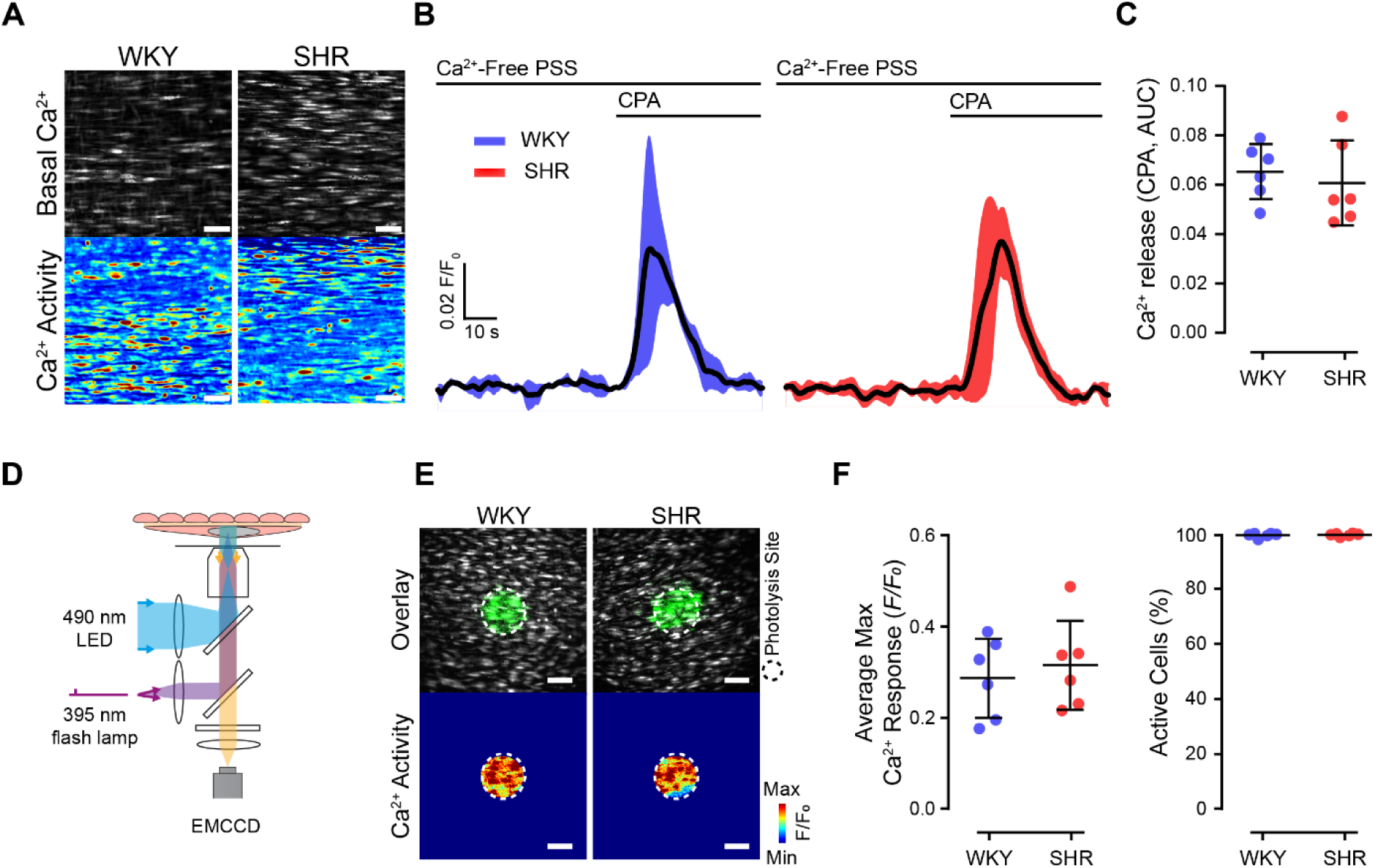
IP_3_ receptor activity and the internal Ca^2+^ store content is maintained in hypertension. A-C) Example Ca^2+^ images (A), mean cyclopiazonic acid (CPA)--induced Ca^2+^ transients (B), and mean ± SD summary data (C) showing the effect of hypertension on the internal Ca^2+^ store content in mesenteric artery endothelial cells. In panel B, the black trace (black line) represents the mean of n = 6 independent trials, the shaded error bands representing the standard deviation of the mean. (D-F) Schematic of imaging system with targeted endothelial cell photoactivation (D), example Ca^2+^ activity images (E), and corresponding mean ± SD summary data (F, n = 6) illustrating the response to activation of endothelial IP_3_ receptors via photolysis of caged IP_3_. Statistical significance (P < 0.05) was assessed using Student’s t-test. Image scale bars = 50 µm.

**Figure S6.**
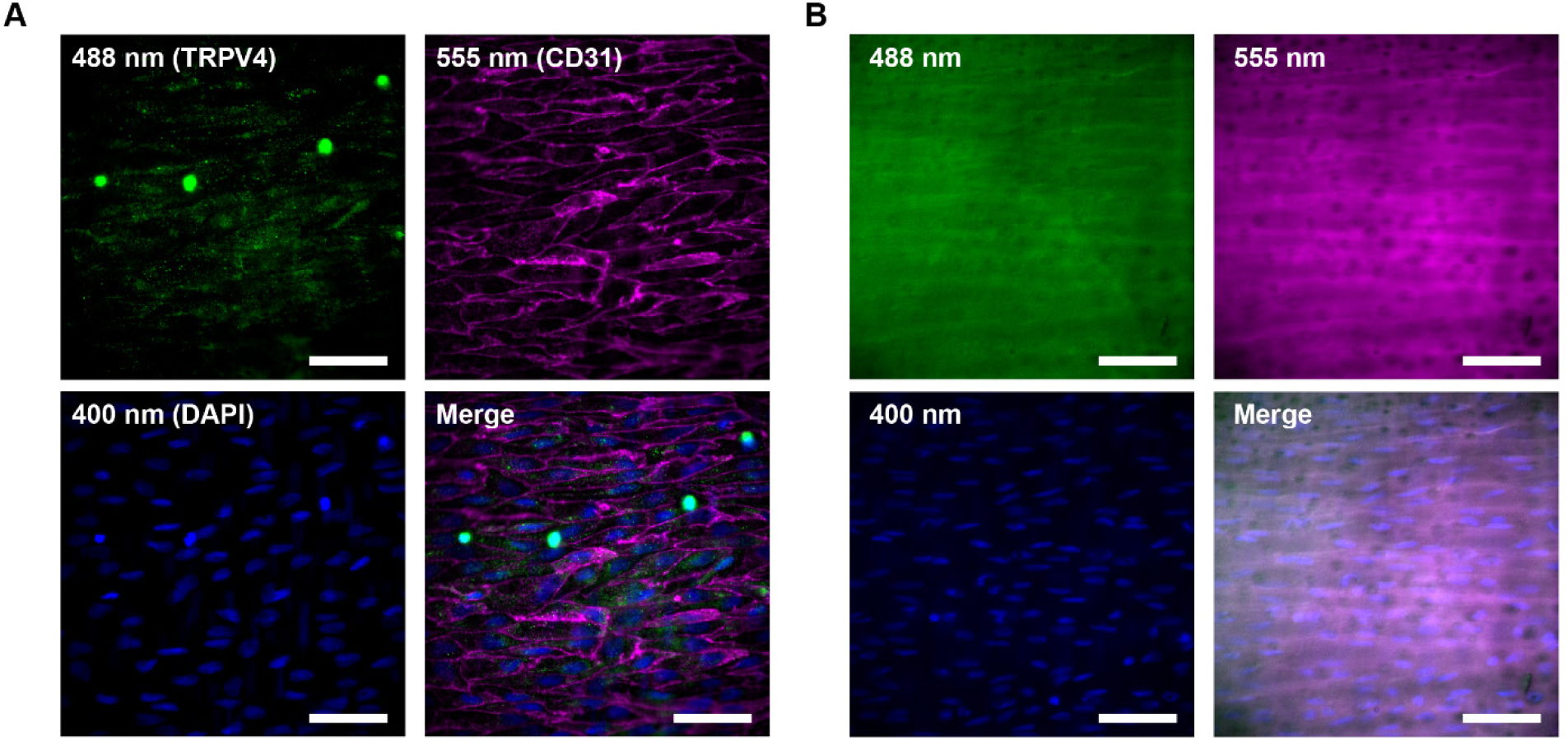
Immunofluorescence staining of endothelial cells in intact arteries. A) Endothelial cell TRPV4 ion channel expression revealed by in situ fluorescence imaging. Images show specific labelling of endothelial cells using primary antibodies against the target antigens: TRPV4 receptor (green), cell-cell borders (CD31, magenta) and nuclei (DAPI, blue). B) Representative images from control experiments with no primary antibody. No fluorescence signal was detected in arteries incubated with secondary antibodies alone. Scale bars = 50 µm.

**Table S1:** Differential Expression Analysis of IP_3_ receptor and TRPV ion channels in Endothelial Cells from mesenteric arteries isolated from WKY and SHR rats.

| Gene | Score | Log Fold Change | P-Value | Adjusted P-Value |
| --- | --- | --- | --- | --- |
| ITPR1 | -6.1 | -1.3 | 2.2E-09 | 0.000 |
| ITPR2 | -0.7 | -0.4 | 5.0E-01 | 1.000 |
| ITPR3 | -2.8 | -1.4 | 5.0E-03 | 0.023 |
| TRPV1 | 0.0 | 0.0 | 1.0E+00 | 1.000 |
| TRPV2 | -0.3 | -0.7 | 7.6E-01 | 1.000 |
| TRPV3 | 0.0 | 0.0 | 1.0E+00 | 1.000 |
| TRPV4 | 2.6 | 0.6 | 1.1E-02 | 0.032 |
| TRPV5 | 0.0 | 0.0 | 1.0E+00 | 1.000 |
| TRPV6 | 0.0 | 0.0 | 1.0E+00 | 1.000 |

**Table S2:** Differential Expression Analysis of IP_3_R and TRPV ion channels in smooth muscle cells from mesenteric arteries isolated from WKY and SHR rats.

| Gene | Score | LogFoldChange | P-Value | Adjusted P-Value |
| --- | --- | --- | --- | --- |
| ITPR1 | -45.7 | -2.6 | 0.0E+00 | 0.000 |
| ITPR2 | 1.4 | 18.7 | 1.6E-01 | 0.472 |
| ITPR3 | -0.6 | -1.2 | 5.2E-01 | 0.891 |
| TRPV1 | -0.5 | -0.4 | 5.9E-01 | 0.891 |
| TRPV2 | 2.1 | 0.4 | 3.5E-02 | 0.157 |
| TRPV3 | 0.0 | 0.0 | 1.0E+00 | 1.000 |
| TRPV4 | -0.7 | -0.3 | 5.1E-01 | 0.891 |
| TRPV5 | 0.0 | 0.0 | 1.0E+00 | 1.000 |
| TRPV6 | 0.0 | 0.0 | 1.0E+00 | 1.000 |

